# Does repeated administration of baclofen in free feeding rats reduce body weight by stimulating brown fat metabolism through activation of the sympathetic nervous system?

**DOI:** 10.1101/2025.01.21.634040

**Authors:** Ivor S Ebenezer

**Affiliations:** Neuropharmacologist Research Group, School of Medicine, Pharmacy and Biomedical Sciences, St Michael’s Building, University of Portsmouth, PO1 2DT, United Kingdom

**Keywords:** GABA, GABA_B_ receptors, Baclofen, Propranolol, Body Weight, Rat

## Abstract

The results of previous research have suggested that chronic administration of the GABA_B_ receptor agonist baclofen can lead to weight loss in rodents by potentially activating brown adipose tissue (BAT) to increase metabolism rate. Studies in anaesthetised rats have shown that baclofen injected directly into the ventromedial nucleus of the hypothalamus (VMH) stimulates BAT to increase thermogenesis. This effect was attenuated by pretreatment with the beta-adrenoceptor antagonist propranolol, suggesting that baclofen microinjected into the VMH stimulate the sympathetic outflow to BAT, consequently increasing metabolic rate. The present study was designed to test this hypothesis by investigating whether pretreatment with propranolol would attenuate the weight loss induced by repeated daily administration of baclofen in free feeding rats. In the first experiment, male Wistar rats were injected intraperitoneal (ip) once daily for 24 days with either saline followed by saline, saline followed by baclofen (4 mg / kg), propranolol (10 or 20 mg / kg) followed by saline, or propranolol (10 mg / kg) followed by baclofen (4 mg / kg), with 5 minutes separating each injection. Body weight was measured each day 24 h after drug administration. Baclofen treatment resulted in significant reductions in daily body weight compared to the saline group. Propranolol alone had no significant effect on body weight. Importantly, pretreatment with propranolol did not attenuate the weight loss induced by baclofen. A similar experimental protocol as described above was used in a second experiment except that the rats were pretreated with a 20 mg / kg dose of propranolol and body weight was measured for 6 days. The high dose of propranolol significantly decreased body weight compared with control data and potentiated the baclofen-induced reduction in body weight. These results suggest that the effects of baclofen of body weight in free feeding rats may not be dependant on the the activation of BAT through stimulation of the sympathetic nervous system (SNS). Further research is therefore necessary to understand how baclofen influences metabolism and weight loss.

## 1. Introduction

The results from a number of studies have shown that while the GABA_B_ receptor agonist baclofen (see Ebenezer, 2015) increases short-term food intake in freely feeding rodents, it has no impact on 24-hour food consumption (Ebenezer and Pringle, 1992, Ebenezer, 1995, Ebenezer, 2012, Ebenezer and Patel, 2004, Higgs and Barber, 2004, Ebenezer and Patel, 2004, Buda-Levin et al., 2005, Patel Ebenezer, 2008a, b, Ebenezer and Patel., 2011., Ebenezer and Prabhaker, 2007, Ebenezer, 2025). However, despite not affecting daily food intake, baclofen administration consistently leads to a dose-dependent reduction in body weight (Ebenezer, 2025, Patel and Ebenezer, 2008b, 2010, Addae et al., 1986). The weight loss has been attributed to an increase in metabolic rate (Addae et al., 1986; Ebenezer and Patel, 2010, Ebenezer, 2025). In support of this suggestion, Addae et al. (1986) demonstrated that chronic administration of baclofen suppressed weight gain in rats by stimulating brown adipose tissue (BAT) activity, thereby increasing metabolic rate. Further, they showed that microinjections of baclofen into the ventromedial nucleus of the hypothalamus (VMH) in urethane anaesthetised rats caused hyperthermia and stimulated BAT thermogenesis. This effect was blocked by propranolol, a beta-adrenoceptor antagonist, suggesting that baclofen acts within the VMH to stimulate the sympathetic outflow to BAT, consequently increasing body temperature and metabolic rate (Rothwell et al., 1985; Addae et al., 1986).

While the studies indicating that weight loss following repeated administration of baclofen in rats may result from central stimulation of sympathetic outflow to BAT (Rothwell et al., 1985; Addae et al., 1986), the relevance of these findings in anaesthetised rats to free feeding animals remains unclear. The aim of the present study was therefore undertaken to determine whether baclofen increases metabolic rate by stimulating the sympathetic nervous system (SNS) by examining the effects of the propranolol on baclofen-induced weight loss in free feeding rats.

## 2. Material and methods

The protocols used in this study were approved by the Ethical Review Committee at the University of Portsmouth, England and carried out under licences granted by the UK Home Office Scientific Procedures Act.

### 2.1 Effects of repeated administration baclofen after pretreatment with propranolol on body weight changes

#### 2.11 Experiment 1

Adult male Wistar rats (n=32; starting body weights: 375 – 455 g; age=16 weeks at start of experiment) were housed in cages in groups of 4 where they had free access to food (food composition: (a) Percentage mass: protein 20%, oil 4.5%, carbohydrate 60%, fibre 5%, ash, 7% +traces of vitamins and metals, (b) percentage energy: protein 27.3%, oil 11.48% and carbohydrate 61.2%, and (c) energy density: 3.6 kcal/g) and water at all times. The animals were maintained on a 12 h light/dark cycle (lights on at 8.30 h and lights off at 20.30 h). During experimental sessions, the rats (n=8 in each group) were injected i.p. once daily for 24 days with either saline followed 5 min later by saline (Group 1a), saline followed by baclofen (4 mg / kg; Group 2a), propranolol (10 mg / kg) followed by saline (Group 3; Group 3a), or propranolol (10 mg / kg) followed by baclofen (4 mg / kg) (Group 4a). Body weight was recorded daily, as described previously (Patel et al., 2010).The body weight data for each rat were expressed as a percentage of the animal’s body weight recorded on the first day of the experiment before administration of saline or drugs

#### 2.12 Experiment 2

In an additional experiment, a similar protocol to that described above for Experiment 1 was used. Male Wistar rats (345 - 415 g, n=6 in each group) were injected i.p. once daily for 6 days with either saline followed 5 min later by saline (Group 1b), saline followed by baclofen (4 mg / kg; Group 2b), propranolol (20 mg / kg) followed by saline (Group 3b), or propranolol (20 mg / kg) followed by baclofen (4 mg / kg) (Group 4b). Body weight was recorded daily, 24 h after saline or drug administration.

### 2.2. Drugs

(±) Baclofen and L-propranolol were purchased from Sigma Biochemicals, Dorset, UK. The drugs was dissolved in physiological saline solution (0.9% w/v, NaCl) to give an injection volume of 0.1 ml / 100 g body weight. Physiological saline solution was used in control experiments.

### 2.3. Statistics

The data was analysed by 2 way analysis of variance (ANOVA) with repeated measures on overall treatment and time (days) followed by the Student-Newman Keul *post-hoc* test (**Winer, 1971**).

### 2.4. Food Intake and Body Weight

The body weight data obtained for each rat were expressed as a percentage of the animal’s body weight recorded on the day the animals received their first injections, denoted as Day 1.

## 3. Results

### 3.1. Effects of repeated administration baclofen after pretreatment with propranolol on body weight changes

(a) Experiment 1. The effects of saline - saline (Group 1a), saline - baclofen (4 mg / kg; Group 2a) propranolol (10 mg / kg) - saline (Group 3a) and propranolol (10 mg / kg) - baclofen (4 mg / kg; Group 4a) on body weight after each feeding session during Days 2 to 25 are shown in Fig. 1. Statistical analysis of the results revealed that there were significant main effects of treatment (F*(3,28)* = 8.71, *P* < 0.01) and time (days) (F*(23,644)* = 42.82, *P* < 0.01) but no significant effects of treatment x time (days) interaction (F*(69,644)* = 1.37, *NS). Post hoc* tests showed that baclofen (Group 2a) significantly reduced body weight gain compared with saline controls (Group 1a) from Day 5 to Day 25 (*P*<0.05 on Days 5 and 6 and *P*<0.01 from Day 7 to Day 25). Propranolol (Group 3a) on its own had no significant effects on body weight changes compared with saline controls. (Group 1a). However, pretreatment with propranolol before administration of baclofen (Group 4a) significantly decreased body weight changes compared with saline controls (Group 1a) from Day 2 to Day 25 (*P*<0.01 on each day). There were no significant differences in body weight changes between the rats in Group 2a (saline -baclofen) and Group 4a (propranolol - baclofen).
(a) Experiment 2. The effects of saline - saline (Group 1b), saline - baclofen (4 mg / kg; Group 2b) propranolol (20 mg / kg) - saline (Group 3b) and propranolol (20 mg / kg) - baclofen (4 mg / kg; Group 4b) on body weight after each feeding session during Days 2 to 7 are shown in Fig. 2. Statistical analysis of the results revealed that there were significant main effects of treatment (F*(3,20)* = 6.96, *P* < 0.01), time (days) (F*(5,100)* = 21.5, *P* < 0.01) and treatment x time (days) interaction (F*(15,100)* = 3.21, *P<*0.01*). Post hoc* tests showed that baclofen (Group 2b) significantly reduced body weight gain compared with saline (Group 1b) from Day 4 to Day 7 (*P*<0.01 on each day). Propranolol (Group 3b) on its own significantly reduced body weight compared to saline controls. (Group 1b) from Day 4 to Day 7 (*P*<0.05 on each day). Pretreatment with propranolol before administration of baclofen (Group 4b) significantly decreased body weight compared with saline controls (Group 1a) from Day 3 to Day 7 (*P*<0.05 on Days 3 and 4 and P<0.01 on Days 5 to 7). The rats (Group 4a) that were pretreated with propranolol (20 mg / kg) prior to baclofen (4 mg / kg) displayed significantly reductions in body weight compared to those that received baclofen after saline (Group 2a) on Days 5 to 7 (*P*<0.05 on each day).

**Figure 1.**
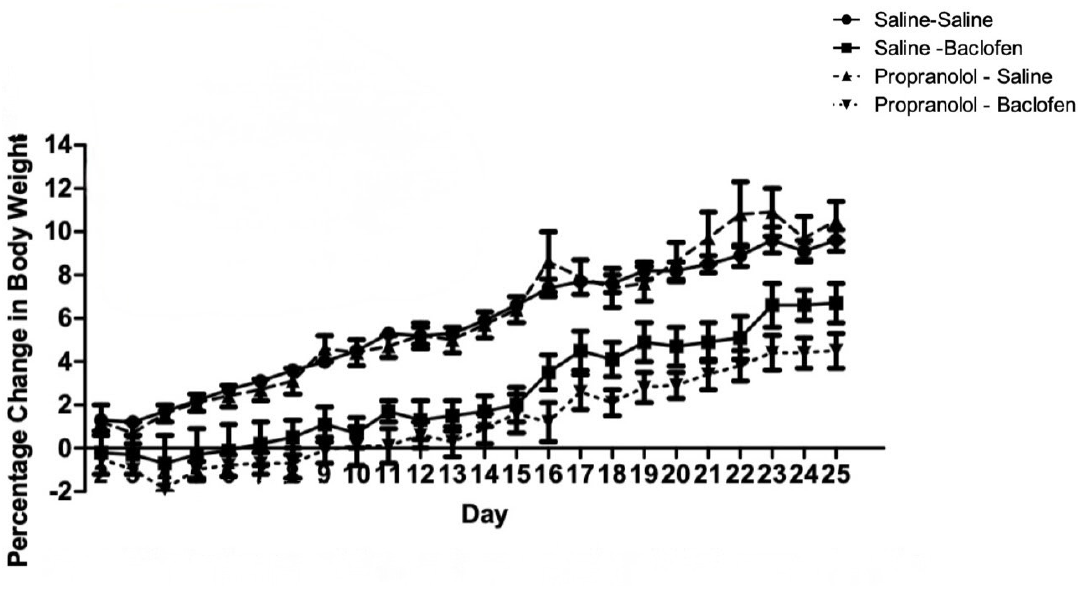
Effects of once daily intraperitoneal injections of physiological saline followed by saline, saline followed by 4 mg / kg baclofen, propranolol (10 mg / kg) followed by saline and propranolol followed by baclofen (4 mg / kg) on percentage changes in body weight from starting weights recorded on experimental Day 1 over a period of 24 days (body weight = 375 – 455 at start of experiment). See text for further details and results of statistical analyses. Vertical lines represent ± S.E.M.

**Figure 2.**
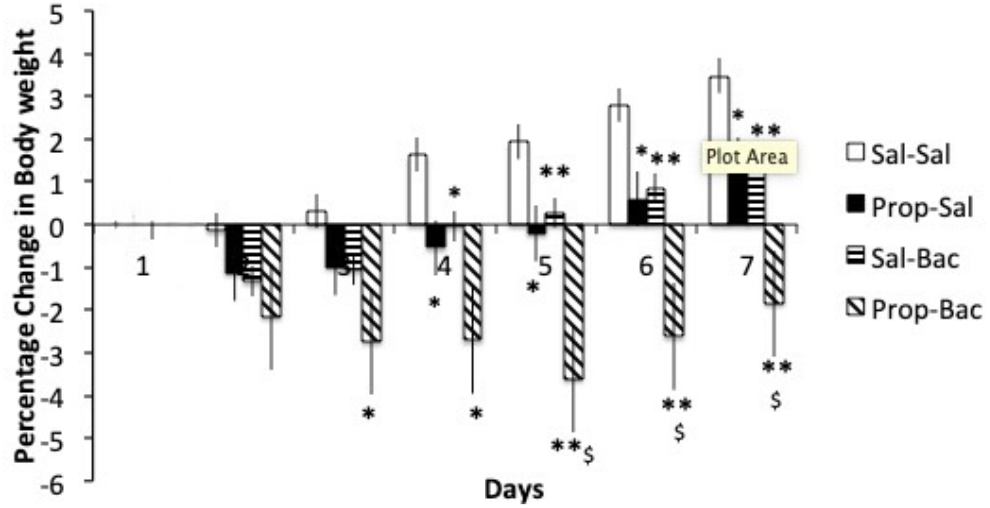
Effects of repeated once daily intraperitoneal injections of physiological saline followed by saline, saline followed by baclofen (4 mg / kg), propranolol (20 mg / kg) followed by saline and propranolol (20 mg / kg) followed by baclofen (4 mg / kg) on percentage changes in body weight recorded on experimental Day 1 over a period of 6 days (body weight = 345 - 415 g at start of experiment). ** P< 0.01 vs saline -saline, *P<0.05 vs saline-saline, ^$^P<0.05 vs saline-baclofen. Vertical lines represent ± S.E.M. See text for further details and results of statistical analyses.

## 4. Discussion

The results from previous studies have indicated that weight loss in rodents observed after repeated baclofen administration may be caused by elevated metabolic rate, as daily food intake remained unaffected (Ebenezer, 2025, Patel and Ebenezer, 2010, Addae et al., 1986). In support of this hypothesis, Addae et al. (1986) found that microinjections of baclofen into the ventromedial nucleus of the hypothalamus (VMH) reduced body weight by activating brown adipose tissue (BAT) metabolism. Importantly, these authors observed that in anaesthetised rats, the increased BAT activity could be inhibited by propranolol, a beta-adrenoceptor antagonist. These findings suggest that baclofen stimulates sympathetic nervous system (SNS) outflow to BAT, thereby increasing metabolic rate. The primary objective of the present study was to extend these observations in anaesthetised rats to free feeding animals by investigating whether pretreatment with propranolol prevents the weight loss following repeated administration of baclofen.

The dose of baclofen employed in this study has been previously demonstrated to reduce body weight in rats (Patel and Ebenezer, 2010). In the first experiment, a 10 mg /kg dose of propranolol was used based on the observation that it decreased 2-hour food intake in fasted rats and the animals did not display overt aversive effects (Ebenezer, unpublished results). Baclofen (4 mg / kg, i.p.) significantly reduced body weight gain in the animals compared to controls, commencing four days after the first injections on Day 1 (Fig. 1). These findings are consistent with previous observations (Patel and Ebenezer, 2010). Propranolol (10 mg / kg, i.p.) alone had no significant impact on body weight compared to saline controls. Moreover, pretreatment with propranolol did not prevent the weight-reducing effects of baclofen, and no significant differences were observed between the saline-baclofen and propranolol-baclofen groups (Fig. 1). These data therefore suggest that daily pretreatment with propranolol (10 mg / kg) does not attenuate the baclofen-induced weight loss by blocking sympathetic activation of BAT as predicted by findings in anaesthetised rats (Addae et al., 1986).

It is possible that the lack of effect of propranolol (10 mg / kg) on baclofen-induced weight loss may have been due to insufficient plasma concentrations of the beta-adrenoceptor blocker. This could be attributed to propranolol’s complex pharmacokinetic profile. There is a paucity of data for the half life of propranolol after i.p. or subcutaneous (s.c.) administration in rats. However, data from intravenous or oral administration in rats have suggested that its estimated half-life is in the range 1.5 to 3 hours, and this may increase due to factors such as age of animal, repeated dosing, increasing doses of drug, strain of rat and route of administration (Iwamoto et al., 1988., Qureshi and Buttar, 2008). Furthermore, the pharmacological actions of the drug may be extended because propranolol is metabolised to 4-hydroxypropranolol, which is an active metabolite with beta-adrenoceptor antagonist properties (Msdubachi et al., 1971). Interestingly, the half-life of baclofen after systemic administration in rats has been estimated to be between 3 to 4.5 hours (Popava et al., 1995, Kim et al., 2014). Given the comparable half-lives of baclofen and propranolol (together with its metabolite) in rats, it is plausible that the 10 mg/kg dose of propranolol would have, at least, partially suppressed sympathetic activity to BAT, assuming baclofen activates it. The potential suppression should have been evident as a reduction in baclofen-induced weight loss. However, this did not occur. A second experiment was therefore undertaken to test whether a higher dose of propranolol (20 mg / kg) would act to reduce the effects of baclofen on body weight.

In contrast to the 10 mg / kg dose, the 20 mg / kg dose of propranolol significantly decreased body weight compared to both the saline and baclofen-treated groups (Fig. 2). The literature on propranolol’s effects on body weight is unfortunately inconsistent, with the data from human and animal studies indicating a range of outcomes. In most cases, there were no effects of body weight but there were some reports that indicated weight gain and, in a few cases, weight loss. One possible explanation for the weight loss observed with the high dose of propranolol is that it may have reduced daily food intake. It has been observed, as mentioned above, that the lower dose of propranolol (10 mg / kg) decreases food intake measured over 2 hours in fasted rats (Ebenezer, unpublished results). However, its long-term effects on feeding are not known. Besides acting as an antagonist at beta adrenoceptors, propranolol also has antagonist activity at central 5-HT_1A_ receptors (Sprouse and Aghajanian, 1986). 5-HT_1A_ agonists, such as 8-hydroxy-2-(di-n-propylamino)tetralin (8-OH-DPAT), increase food intake in rats (Ebenezer, 1992). Therefore, blocking these receptors could theoretically lead to a reduction in food intake. However, this is unlikely as WAY 100635, which is a potent 5-HT_1A_ receptor antagonist, does not decrease food consumption when administered on its own (Ebenezer et al., 2007). Another possibility is that the high dose of propranolol acts centrally or exerts effects on organs, tissue and glands innervated by the SNS to affect ingestive behaviours. This could lead to weight loss through several mechanisms. For example, propranolol may stimulate or inhibitthe release of hormones involved in regulation of feeding, it could interact with central or peripheral neurotransmitters and neurochemicals involved in regulating appetite or the high dose might elicit non-specific aversive effects in the animals leading to reduced food intake. The observation that 20 mg / kg propranolol not only reduced body weight on its own but significantly enhanced the weight-suppressing effect of baclofen (Fig.2) indicates that, contrary to findings in anaesthetised rats (Addae et al., 1986), it does not attenuate baclofen-induced weight loss by inhibiting sympathetic activation of BAT.

The results of this study therefore suggests that the reduction in body weight observed with baclofen may not primarily stem from increased SNS stimulation to BAT. While baclofen may activate thermogenesis to increase metabolic rate, its mechanism of action might involve direct effects on BAT itself. Supporting this suggestion, GABA_B_ receptors have been identified on BAT (see Ikegami et al., 2018), suggesting a potential for direct activation of BAT by baclofen. Furthermore, repeated baclofen administration could induce “browning” of white fat, a process where white fat cells acquire characteristics of brown fat, enhancing their metabolic activity and calorie burning characteristics (Machado et al., 2022). This “browning” can be triggered by various factors, including cold exposure, exercise, and certain nutrients, such as omega-3 fatty acids (see Machado et al., 2022). Interestingly, we have found that repeated daily administration of baclofen to free feeding rats increases browning of white fat (Bains and Ebenezer, unpublished results). Therefore, the observed weight loss with baclofen may result from a combination of direct effects on BAT and the induction of white fat browning.

In conclusion, the results of this study suggest that baclofen’s effects on body weight in free feeding rats may not be primarily mediated by activation of BAT through sympathetic nerve stimulation. Further research is therefore needed to elucidate the precise mechanisms by which baclofen influences metabolism and induces weight loss.

## References

Addae, J.I., Rothwell, N.J., Stock, M.J., Stone, T.W., 1986. Activation of thermogenesis of brown fat by baclofen. Neuropharmacol. 25, 627–631.

Buda-Levin, A., Wojnicki, F.H.E., Corwin, R.L., 2005. Baclofen reduced fat intake under binge-type conditions. Physiol. Behav., 86, 176–184.

Ebenezer, I.S., 1992. Effects of the 5HT_1A_ agonist, 8-OH-DPAT, on operant food intake in non-deprived rats. NeuroReport, 3, 62–64.

Ebenezer, I.S., 1995. Intraperitoneal administration of baclofen increases consumption of both solid and liquid diets in rats. Eur. J. Pharmacol. 273, 183–185.

Ebenezer, I.S., 2015. Neuropsychopharmacology and Therapeutics, John Wiley & Sons, Chichester, UK, ISBN 978-1-118-38565-4

Ebenezer, I.S., 2023. The effects of repeated intraperitoneal administration of the GABA_B_ receptor agonist baclofen on body weight in rats maintained on a restricted feeding schedule. BioRxiv, 10.1101/2023.01.24.525410

Ebenezer, I. S., 2025. Repeated intraperitonral administration of the GABA_B_ receptor agonist baclofen reduces body weight in the mouse. Biorxiv, 10.1101/2025.01.05.631370

Ebenezer, I. S., Arkle, ML., Tite, R.M. 2007. 8-Hydroxy-2-(di-n-propylamino)tetralin (8-OH-DPAT) inhibits food intake in fasted rats by an action at 5-HT_1A_ receptors.Meth. Find, Expt. Clinton. Pharmacol. 29, 269 – 274.

Ebenezer, I.S., Patel, S.M., 2004. Effects of the GABA_B_ receptor agonists baclofen and 3-aminopropylphosphinic acid (3-APA) on food intake in rats. Meths. Find. Expt. Clin. Pharmacol. 26, 627–630.

Ebenezer, I.S., Patel, S.M., 2011. Effects of intraperitoneal administration of the GABA_B_ receptor agonist baclofen on food intake measured under different feeding conditions. Eur. J. Pharmacol.653,58–62.

Ebenezer, I.S., Pringle, A.K., 1992. The effects of systemic administration of baclofen on food intake in rats. Neuropharmacol.31, 39–42.

Ebenezer, I.S., Prabhaker, M., 2007. The effects of intraperitoneal administration of the GABA_B_ receptor agonist baclofen on food intake in CFLP and C57BL/6 mice. Eur. J. Pharmacol. 569,90 – 93.

Higgs, S.., Barber, D.J., 2004. Effects of baclofen on feeding behaviour examined in the runway. Prog. Neuro-Psychopharmacol. Biol. Psychiat.28, 405 – 408.

Machado, SA., Nascimento, G. P., da Silva, D.S., et al., 2022. Browning of the white adipose tissue regulation: new insights into nutritional and metabolic relevance in health and disease. Nutrition &Metabolism, 19, 61. 10.1186/s12986-022-00694-0

Patel, S., Ebenezer, I.S., 2008a. The effects of chronic intraperitoneal administration of the GABA_B_ receptor agonist baclofen on food intake in rats. Eur. J. Pharmacol.593, 68–72

Ikegami R., Shimizu, I, Sato, T., et al., 2018. Gamma-aminobutyric acid signalling in brown adipose tissue promotes systemic metabolic derangement in obesity. Cell Reports, 24, 2827 – 2827.

Iwamoto, K., Watanabe, J., Aoyama, Y., 1988. Age-dependent pulmonary first-pass elimination of propranolol in rats. J.Pharm. Pharmacol. 40, 135 – 137.

Kim, T.H., Park, G-Y.m Shin, S., et al., 2014. Evidence Based Complementary and Alternative Medicine. 10.1155/2014/402126

Masubachi, Y., Hosokawa, S., Horie, T., et al., 1971. Pharmacology of 4-hydroxypropranolol, a metabolite of propranolol. Br. J. Pharmacol. 43, 222 – 235.

Patel, S., Ebenezer, I.S., 2008b. The effects of acute multiple intraperitoneal injections of the GABA_B_ receptor agonist baclofen on food intake in rats. Eur. J. Pharmacol. 601, 106–110.

Patel, S.M., Ebenezer, I.S., 2010. Effects of chronic administration of the GABA_B_ receptor agonist baclofen on food intake and body weight in rats. Eur. J. Pharmacol.635, 129–134.

Popova, E.D., Puzin, M.N., Kolyvanov, G.B., Levin, A.A., 1995. Baclofen pharmacokinetics in rats. Eksp. Klin. Farmakol. 58, 53–54.

Qureshi, S.A., Buttar, H.S., 1988. A comparative study of the pharmacokinetics of propranolol and its major metabolites in the rat after oral and vaginal administration. Xenobiotica. 19, 833. - 890.

Rothwell, N.J., Addae, J.L., Stock, M.J., Stone, T.W., 1985. Baclofen is a potent activator of brown fat metabolism. J.Pharm.Pharmacol. 37, 926 – 927.

Sprouse, J.S., Ahajanian, G.K., 1986. (-) Propranolol blocks the inhibition of serotonergic dorsal raphe firing by 5-HT_1A_ selective agonists. Eur. J. Pharmacol., 128, 295 – 298.

Winer, B.J.. 1971. Statistical principles in experimental design. New York, McGraw-Hill.

